# Influences of single mutation and retinal binding on the THz absorption spectra of CRABP-II based rhodopsin mimics

**DOI:** 10.1101/2024.04.28.591535

**Authors:** Yunyu Wang, Yongnan Hu, Jiajia Meng, Xubiao Peng, Qing Zhao

## Abstract

The collective vibration of many biomolecules such as the skeleton vibration, dipole rotation and conformational bending falls in the terahertz (THz) frequency domain. Terahertz time-domain spectroscopy (THZ-TDS), which is very sensitive to the conformational changes, can be used to characterize the collective vibration of biomolecules. In this study, we investigated the low-frequency THz absorption spectra of two rhodopsin mimics using transmission THz-TDS. Using the normal model analysis (NMA), we successfully modelled the experimental terahertz absorption curve and attributed a unique collective motion pattern to each distinctive terahertz absorption frequency. By comparing the terahertz absorption spectra between without and with retinal, we show that the retinal binding can significantly alters the terahertz absorption spectra as well as the vibration modes. Furthermore, by comparing the terahertz absorption spectra between the two mutants, we observed that the single mutation can significantly change the influence of retinal binding on the terahertz absorption spectrum.

## Introduction

Terahertz (THz) radiation falls in the electromagnetic spectrum, occupying the frequency range between microwaves and infrared, specifically spanning from 0.1 to 10 terahertz (THz)^[1,2]^. THz-TDS is a low-energy coherent detection technique that effectively extracts a range of optical parameters, encompassing refractive index, absorption coefficient, and dielectric constant, et al, by recording the amplitude and phase information of the transmitted terahertz electric field^[3,4]^. Terahertz spectrum contains the chemical information of the molecular conformation that is closely related to the functions of the biological macromolecules, so the terahertz absorption spectrum is also called biological fingerprint spectrum ^[5]^. Simultaneously, due to the inherently low energy of terahertz, it avoids damaging the molecular structure of the sample during the detection process, thereby preserving the integrity of the sample information^[6]^.

In recent years, more and more researchers used THz-TDS as a new technology to reveal the weak interaction between biomolecules, molecular conformation, and its dynamics^[7,8,9]^. In particular, Markelz et al. used DNA, bovine serum protein, and collagen to demonstrate the optically active low-frequency vibration patterns of proteins in the terahertz frequency range^[10]^.

As a photosensitive protein, rhodopsin plays an important role in the life of animals and microbial organisms. It can absorb visible light across various wavelengths and carry out crucial biological functions like signal transduction^[11]^. Hence, understanding the structural and dynamical properties of the rhodopsin is important. Several studies have been conducted on the terahertz absorption of rhodopsin to elucidate its dynamic behavior. KN Woods employed THz-TDS to explore various conformations of bacterial rhodopsin (BR)^[12]^. Additionally, R. Balu et al. investigated the low-frequency terahertz spectra of two photoactive proteins, rhodopsin and bacterial rhodopsin, to characterize the collective low-frequency motion of helical transmembrane proteins. This research underscores the significance of terahertz spectroscopy in identifying vibrational degrees of freedom^[13]^. Furthermore, the work of Groma *et al*. revealed the fundamental primary charge shift phenomenon occurring during the functional energy conversion of bacterial rhodopsin (BR)^[14]^. Dong-Kyu Lee *et al*. observed the light-induced conformational changes in rhodopsin using ultrafast terahertz molecular sensors, suggesting that terahertz absorption spectra can be used to characterize the conformational changes in rhodopsin^[15]^.

Nonetheless, due to the intricate nature of natural rhodopsin as a membrane protein containing over 200 residues, studying such a sophisticated system can be challenging^[16]^. Therefore, researchers have devised various rhodopsin mimics, which exhibit remarkable stability across mutations without requiring membrane integration. This advancement has facilitated the investigation into the wavelength modulation mechanism of rhodopsin^[17]^. In this study, we employed broadband terahertz time-domain spectroscopy, spanning from 0 to 2.5 THz, to explore the terahertz absorption spectra of two CRABPII-based rhodopsin mimics that differ by a single residue. Specifically, we focused on the mutants M2 and M9, designed by Meisam Nosrati et al^[18]^, which we relabeled as R1 and R2 for convenience. The sequence alignment of R1 and R2 is shown in **Figure S1**, where we can see that R2 differs from R1 in a single mutation A32W. We note that researchers mostly focused on the absorption response of protein within 0.1∼1.5 THz in previous studies ^[12]^. Hence, the frequency range (0.1∼2.5THz) considered in our study is an extension of previous researches.

In addition, previous studies have shown that the presence of water in protein solutions will cause a monotonic increase in absorption with frequency, thus concealing the distinct spectral signatures specific to the protein itself^[19]^. This remains a significant challenge in the terahertz absorption spectra of proteins. In this study, we used the freeze-dried samples to effectively eliminate the influence of water molecules, which enabled us to reveal the characteristic terahertz spectrum of the protein more clearly^[20]^. Although the dynamics of the freeze-dried protein may differ from its behavior in a physiological environment, this system nevertheless offers a valuable representation of the protein’s behavior in a vacuum state, thereby enabling quantitative analysis using the normal mode analysis (NMA) method.

Using NMA, we investigated the correlation between the observed terahertz absorption spectra and the vibration modes of these rhodopsin mimics. Additionally, we examined how retinal binding affects the terahertz spectra and vibration modes of both mutants. Our comparative analysis revealed significant differences in the changes of terahertz spectra induced by retinal binding between the two mutants, emphasizing the substantial impact of this solitary mutation.

## Materials and methods

### 1: Sample’s preparation

In our study, the liquid protein samples of R1 and R2 are performed in the laboratory from plasmid extraction, strain culture, and protein purification. Rhodopsin mimics are expressed by Escherichia coli, purified by a His-Tagged nickel column, and detected by electrophoresis **(Figure S2b)**. The protein is centrifuged and concentrated to 15mg/mL and stored in Tris-HCl buffer (pH 7.8), where part of the sample is bound to the protein ligand retinal in a molar concentration ratio of 2:1. In this article, the proteins bound to retinal are labeled as “R1 + retinal” and :R2 + retinal”. The liquid sample is vacuum lyophilized to eliminate the influence of water molecules and then ground into powders, which are mixed with a certain proportion of polytetrafluoroethylene (PTFE) and pressed into round flakes with a 1.3-mm diameter and a thickness gradient **(Figure S2c)**. In the process of grinding and compression, it is necessary to pay attention to the ratio between protein and diluent PTFE and the thickness of the tableted sample, so as to keep the THz transmission signal neither too strong nor too weak in the following THz-TDS experiment. In this study, the mass fraction of 1:9 between protein and PTFE is used. By dissolving the tableted sample in the buffer and observing the color changes resulting from the protein’s photoisomerization, we verified that the protein remained intact despite the compression.

### 2: Transmission terahertz time-domain spectroscopy

In our study, the THz time-domain system came from Beijing Daheng Optoelectronics company. The schematic diagram of THz-TDS system in transmission mode is shown in **Figure 1**. The femtosecond laser pulse produced by the sapphire oscillator has a maximum duration of 50 fs, a central wavelength of 800nm, a repetition frequency of 800 MHz, and an average power of the detected beam greater than 400mW. In the transmission spectrum, the spectrum bandwidth provided by the system is 0.1∼3.5 THz, the dynamic range is 60dB, the sampling interval of THz-TDS is femtosecond, the spectral resolution is 10GHz, and the beamwidth diameter of the terahertz time-domain imaging system at the focal spot is about 5mm. The terahertz time domain system used was enclosed in a dry nitrogen purification tank, which was kept below 5% system humidity during experiments to reduce terahertz absorption due to ambient humidity.

**Figure 1:**
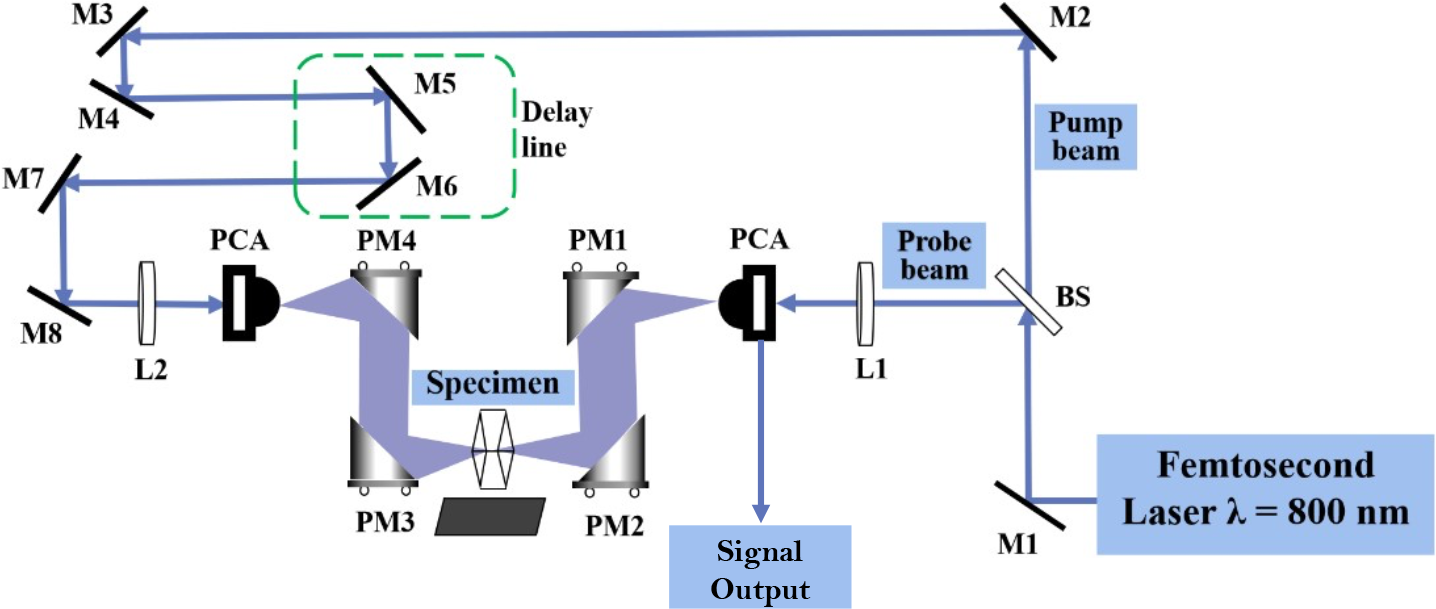
Schematic diagram of transmissive terahertz time-domain spectral path. (M1-M8: Reflector; PCA: photoconductive antenna; BS: Beam splitter.)

### 3: Calculation of refractive index and absorption coefficient

The formula for calculating the refractive index (Eq.(1)) and the expression for the absorption coefficient (Eq.(2)) of the sample are derived from the Beer–Lambert law of attenuation of Fresnel reflection and refraction law. The detailed derivation process can be seen at **Supplementary information**.

The refractive index and absorption coefficient of the sample can be expressed as^[21]^:

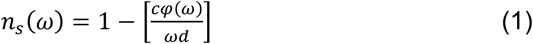

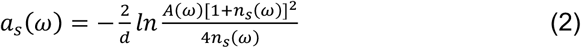

where, *n*_*s*_(*ω*) is the refractive index of the sample, *a*_*s*_ (*ω*)is the absorption coefficient of the sample in the terahertz band. Φ(ω) is the phase difference between the sample signal and the nitrogen transmission signal after Fourier transformation. *c* is the propagation speed of THz wave in nitrogen environment, ω is the angular frequency, and *d* is the sample thickness.

### 4: Calculation of the terahertz spectra with NMA

Normal mode analysis (NMA) assumes that the lowest frequency mode is caused by the maximum movement of proteins and is suitable for studying low-frequency patterns in large molecules such as proteins in vacuum or solid state^[22]^.

We followed the method provided by the Adam J. Mott and David van der Spoel to calculate the terahertz absorption spectra based on the eigenfrequencies and eigenvectors obtained from NMA^[23,24,25]^. The structure is first energy minimized in double precision with force field CHARMM 36m, followed by the NMA to obtain the vibrational eigenfrequency and eigenvector. Both procedures were performed by software GROMACS-2018.6. In the energy minimization process, we took the conjugate gradient algorithm. The initial structure for energy minimization on R1 was directly taken from X-ray crystallographic structure in Protein Data Bank (PDB) (PDBID: 4YFQ), while for R2 it was obtained by mutating the residue A32 to tryptophan in PDB 4YFQ according to the rotamer library ^[26]^ using the software UCSF Chimera^[27]^. The energy minimized structures of R1 and R2 are superimposed in **Figure 2**, where we can see that mutants R1 and R2 have similar overall structures and the greatest deviation between them is located in the α-helix nearby the chromophore retinal.

**Figure 2:**
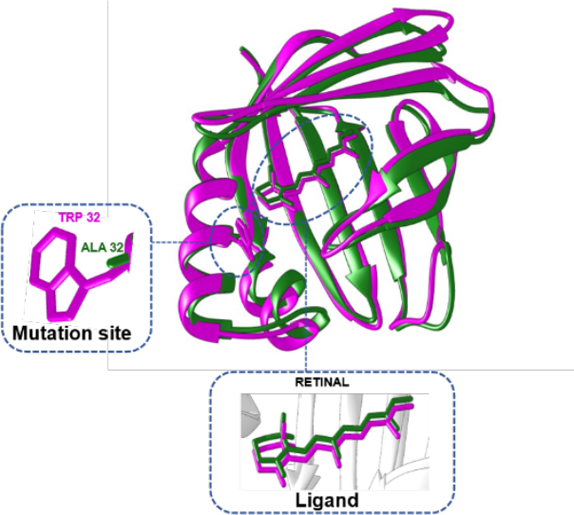
The comparison of the energy minimized structures of R1 and R2. The purple part is the structure of protein R2, whereas the green part is the structure of protein R1. The mutation site and the ligand retinal are depicted in stick representation, with detailed views provided in the zoomed-in insets.

After NMA the terahertz absorption intensity was obtained by equation

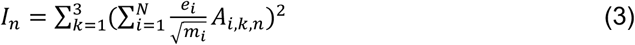

where *I*_*n*_ is the intensity of the *n* th vibrational mode, *k* is the direction index (the x,y,z dimension), *i* is the index of atom (from 1 to *N*), *e*_*j*_ is atomic charge of the *i* th atom, *m*_*i*_ is the mass of atom, and *A*_*i,k,n*_ is element of the *n* th normal mode eigenvector (*N*-by-3 matrix) corresponding to the *i*th atom and in direction *k*. For the derivation details of Eq (3), we refer to the references [24-26]. Since our experiment only measured the lower terahertz region, our calculations are truncated at the 2THz.

For each eigenfrequency, the trajectory of the protein vibration mode can be obtained from the corresponding eigenvector, from which the root mean square fluctuation (RMSF) was then calculated. Finally, the RMSF is normalized through dividing it by its maximal value in order to compare the most fluctuated regions under different eigenfrequencies.

## Results and discussions

### 1: Terahertz absorption spectrum

#### 1. THz-TDS of proteins

Three groups of protein tablets with different thicknesses can be obtained during sample preparation. The thickness gradients of the tabled samples are 0.5 mm, 0.65 mm, and 0.8 mm, respectively, regardless of the binding of the R1 or R2 protein to retinal **(Figure S3)**. The protein without retinal is a white round flake, and the lyophilized sample binding to retinal is a pale yellow round flake due to the dissociation of the retinal part during the lyophilization or tableting process. Terahertz time-domain spectroscopy was measured for each of these samples, and terahertz absorption spectra were characterized in the range of 0.1–2.5 THz.

Firstly, we measured the THz wave directly passing through nitrogen environment as the reference signal in our experiments. The THz signal passing through PTFE was also measured as shown in **Figure S4**. We can see that the PTFE has almost no absorption in THz signals, so it is safe to be used as diluent. We then kept the experimental environments unchanged, such as the temperature T, humidity, etc., and measured the transmittance time-domain signal of the THz passing through the sample, as the sample signal.

The influence of retinal binding on THz absorption differs between R1 and R2, as evident in **Figures 3a–3b** at a 0.8-mm thickness as an example. Specifically, terahertz absorption increases when the retinal is binding to R1, whereas it decreases in R2. The occurrence of THz time-domain peaks is delayed as the thickness of the sample increases. This delay is due to the propagation time required within the material. **Figure 3c** clearly demonstrates that there is a linear correlation between the timing of spectral peaks in the time domain and the compression thickness of various samples. Additionally, we observe that as the sample thickness uniformly increases, there is a corresponding and relatively uniform decrease in the primary peak of terahertz transmission **(Figure 3d)**. This finding indicates that the absorption strength of the THz signal increases with thicker samples, primarily because of the augmented protein content. Hence, this study primarily focused on analyzing the THz absorption spectra of the 8-mm thick samples. Additional experimental data can be accessed in the supplementary information for further reference.

**Figure 3:**
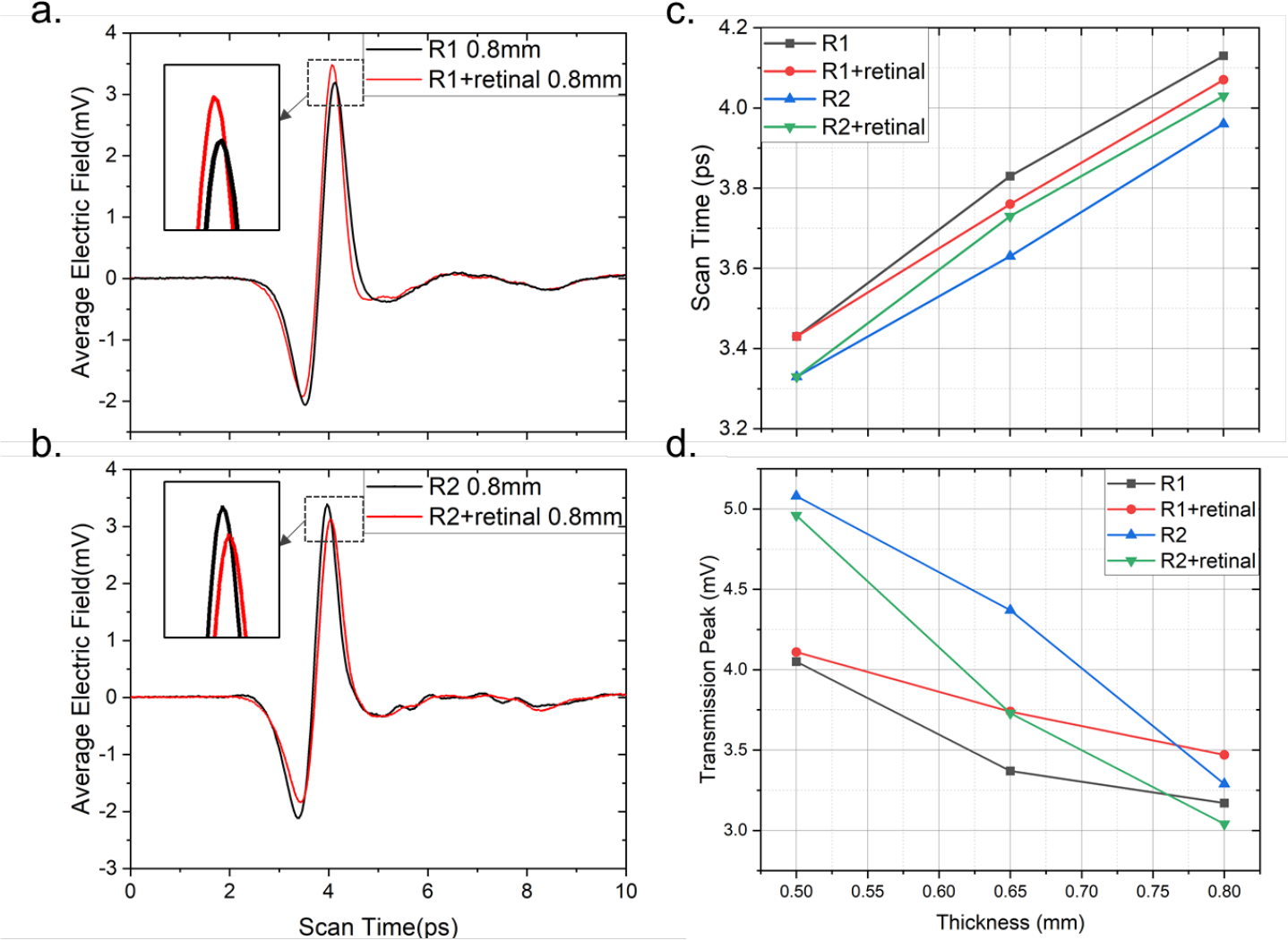
Terahertz transmission time-domain spectra of protein samples of different thicknesses. a: THz-TDS of R1 and R1+retinal. b: THz-TDS of R2 and R2+retinal protein. c: The change in the spectral peak in the time domain with thickness. The “Scan Time” represents the timepoints when the peak occurs. d: The peak value of protein transmission varies with thickness in all four cases

### 2. Terahertz frequency domain spectra

The terahertz time-domain signals were converted into the frequency-domain transmission spectra for both the reference (the nitrogen transmission signal) and the rhodopsin mimic sample through Fourier transformation. Then we obtained the refraction index of the samples from **Equation (1)**, as presented in **(Figure S5)**. From **Equation (2)**, we further obtained the absorption spectra spanning the 0.1-2 THz range, shown as the black lines in **Figure 4**. We also calculated the corresponding absorption spectra from the NMA method as described in Method section, shown as the blue lines and blue bars in **Figure 4**. The blue bars are the primary absorption intensities calculated by **Equation (3)** at the eigenfrequencies from the NMA (only the higher intensities shown), and the blue curve is the theoretical terahertz absorption spectra obtained from addition of all the blue bars with Lorentz broadening with full width half maximum (FWHM) 2cm.

**Figure 4:**
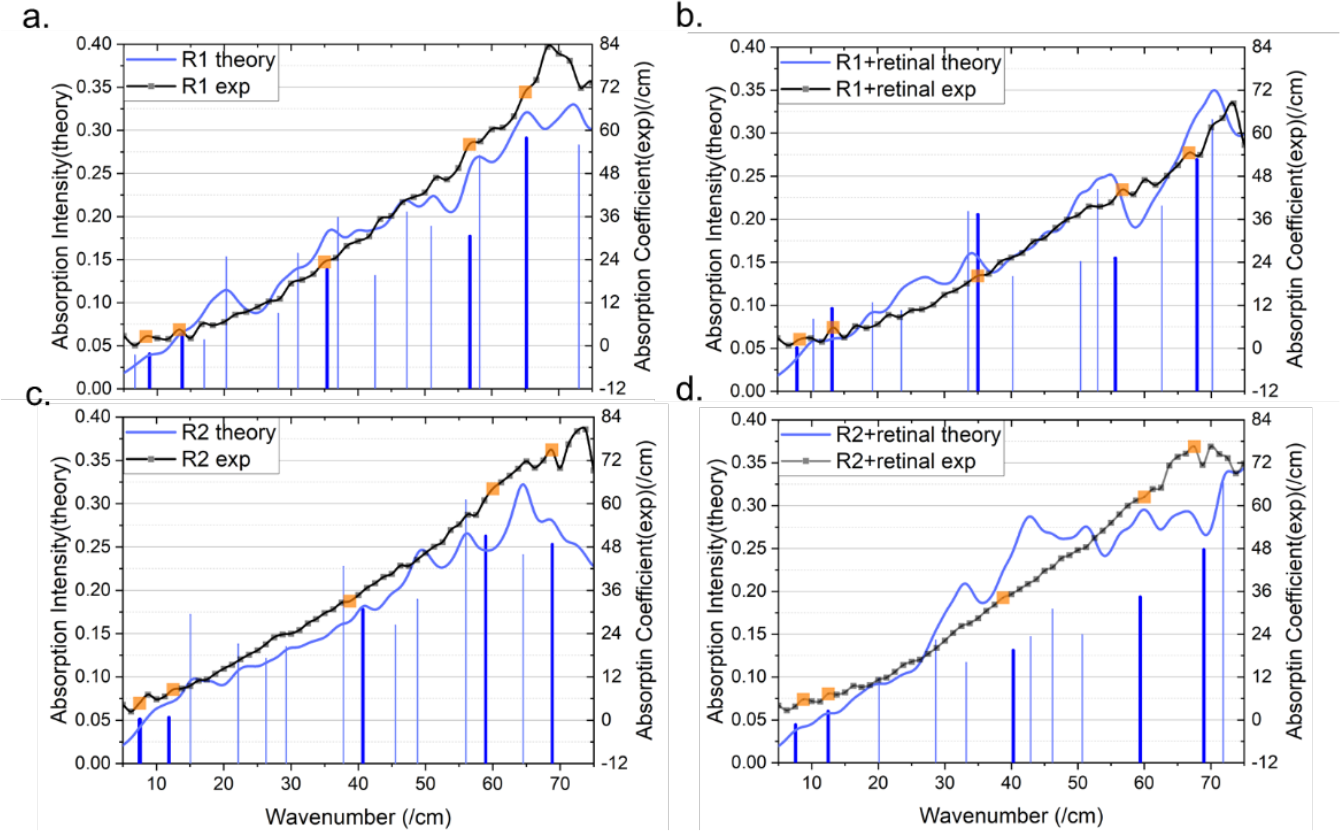
Theoretical and experimental terahertz absorption spectra of proteins. The solid black lines represent the experimental spectrum. The solid blue lines represent the theoretical spectrum. The blue bars represent the theoretically calculated terahertz absorption eigenfrequencies, of which the thicker blue bars will be discussed in detail later. The orange dots are the experimental absorption peaks that fit well with the theory.

From **Figure 4**, we can see that the terahertz absorption spectra exhibit an overall trend of increasing as the frequency rises within the frequency domain 0∼75 cm^−1^. Such an observation is consistent with previous studies on the bacterial rhodopsin and animal rhodopsin ^[13]^, where the frequency range was limited to 0∼1.5THz. In addition, the terahertz absorption spectra presented in Figure 4 exhibit several local small oscillations, reflecting the more precise absorption peaks associated with specific vibration modes of the protein.

A comparison of the calculated absorption spectra with experimental results reveals a strong alignment between the two. The calculated spectra not only follow the overall trend of the experimental data but also exhibit good agreement with the local small oscillations, particularly in the case of absence of retinal. In the case of retinal binding, there is a significant deviation between theoretical calculation and experimental results. One possible explanation for this deviation is that not all retinal molecules successfully bound to the proteins in our sample, resulting in minor quantities of unbound retinal and proteins remaining. This residual material introduces noise into the terahertz absorption spectra, potentially affecting the accuracy of our measurements.

Despite this deviation, both experimental and theoretical results arrive at similar conclusions when comparing the spectra of the protein without retinal to that of the protein with retinal bound. Notably, when the protein binds to retinal, the experimental absorption spectra exhibit fewer local small oscillations and a smoother curve. Correspondingly, our theoretical calculation also indicates a reduction in eigenfrequencies following retinal binding.

In particular, we have identified several eigenfrequencies that closely align with the peaks of small oscillations observed in the experimental absorption spectra, where the theoretical eigenfrequencies are labelled in thicker bars, while the corresponding experimental frequencies are highlighted with bright large squares in Figure 4. In the subsequent sections, we will delve into a detailed discussion on the vibration modes associated with these eigenfrequencies.

### 2: Vibration modes through normal mode analysis (NMA)

We obtained the trajectory with 30 frames of the vibration mode from the corresponding eigenvector for each calculated eigenfrequency in Figure 4. We then calculate the RMSF of each residue from the trajectory to show the flexibility of each residue along the protein^[28]^. The residues with large RMSF are those vibrating ones. Since the overall vibration amplitude can vary significantly at different eigenfrequencies, we normalized the RMSF values for each eigenfrequency, as outlined in the Method section. This normalization allows us to compare the vibration regions across various eigenfrequencies. In Figure 5, we show the normalized RMSF for proteins R1 and R2 without and with binding to retinal at all the eigenfrequencies, respectively. To quantitatively identify the vibration regions, we establish a threshold value of 0.9. Specifically, for each eigenfrequency, only the regions where the normalized RMSF exceeds this threshold will be designated as the vibration regions.

**Figure 5:**
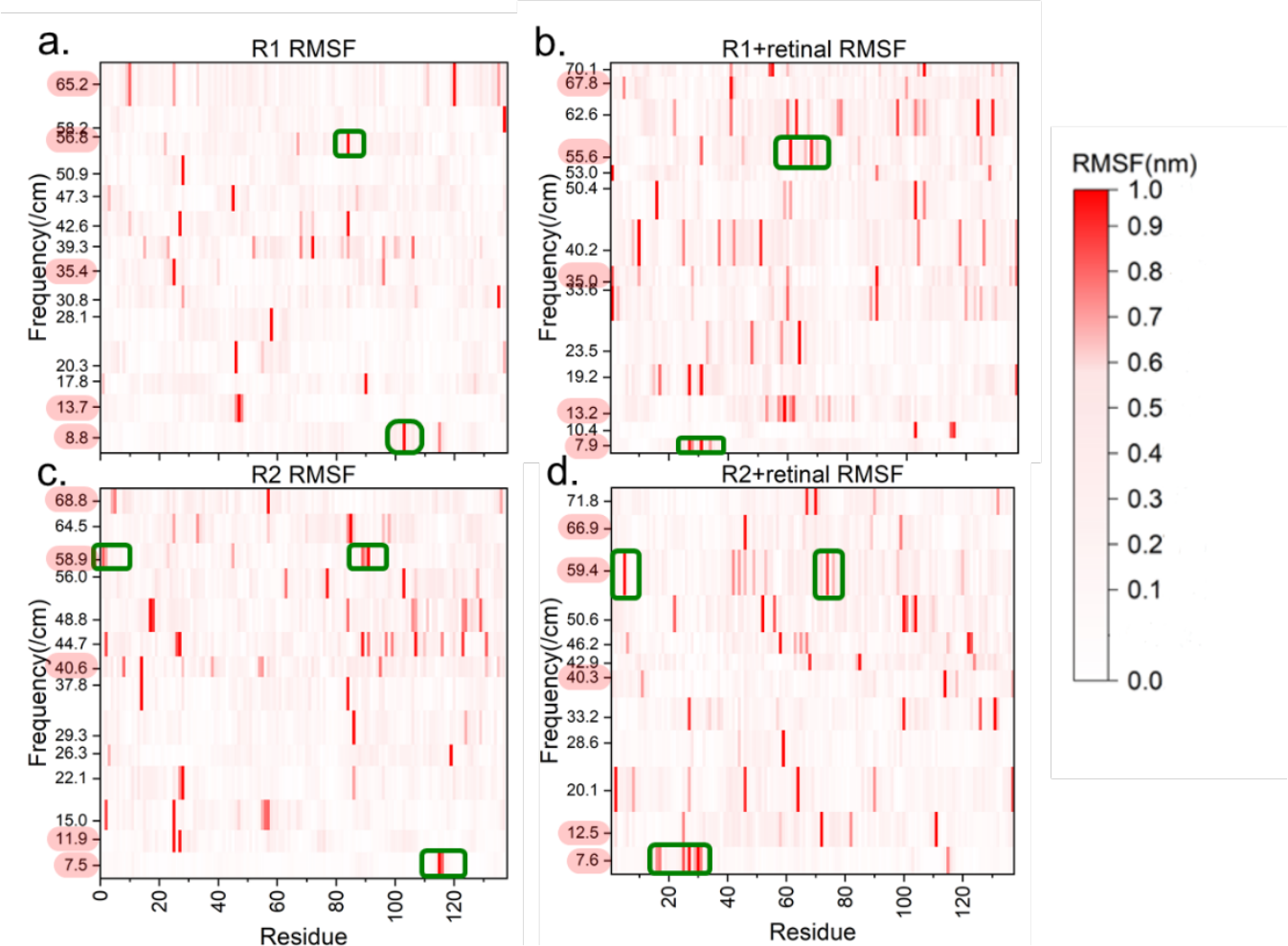
RMSF heat map of rhodopsin mimics. The x-axis represents the residue index, the longitudinal axis represents the terahertz absorption eigenfrequency, and the color shade represents the value of RMSF.

We observe that retinal binding increases the vibration regions corresponding to similar eigenfrequencies by comparing the RMSF maps of the system with and without retinal binding in Figure 5. In particular, the vibration regions of R1 and R2 are both located in the range of residue index 110–120, with relatively sparse vibration residues, at the lowest eigenfrequencies. However, upon the binding of retinal, the vibration area shifts to the range of residue index 20∼40, and the vibration residues become significantly dense. Similarly, at the high eigenfrequency of around 58cm^−1^, the vibration region for R1 is confined to the residue index range of 70∼90, while in R2, it spreads out, encompassing regions in the N-terminal and around residue index 90. With retinal binding, the vibration regions become even more widespread, as evident from the comparison between **Figures 5a/5c** and **5b/5d**. These changes are clearly highlighted in **Figure 5**, where the vibration regions are labeled with green boxes. Such an observation indicates that the retinal binding leads to a more extensive and complex pattern of vibrations mode due to the interaction between protein and retinal.

In addition, a comparison between **Figure 5a** and **Figure 5c** reveals that mutation A32W does not significantly impact the vibration mode in the low-frequency range (0∼30cm^−1^). However, in the mid-to-high-frequency range (30∼70cm^−1^), the vibrating region in R2 is larger than in R1 at the similar eigenfrequencies. We speculate that this is due to the fact that the tryptophan is a hydrophobic residue with long sidechains, leading to the extra interactions with other residues of the protein.

The several eigenfrequencies selected in **Figure 4**, which exhibit good agreement with the experimental results, are indicated by pink shade along the y-axis in **Figure 5**. Additionally, **Figure 6** presents the corresponding 3D structures, emphasizing the vibration regions in the representations of sphere. For the vibrations mode in term of cartoon format showing the vibration directions, we refer to **Figure S6**.

**Figure 6:**
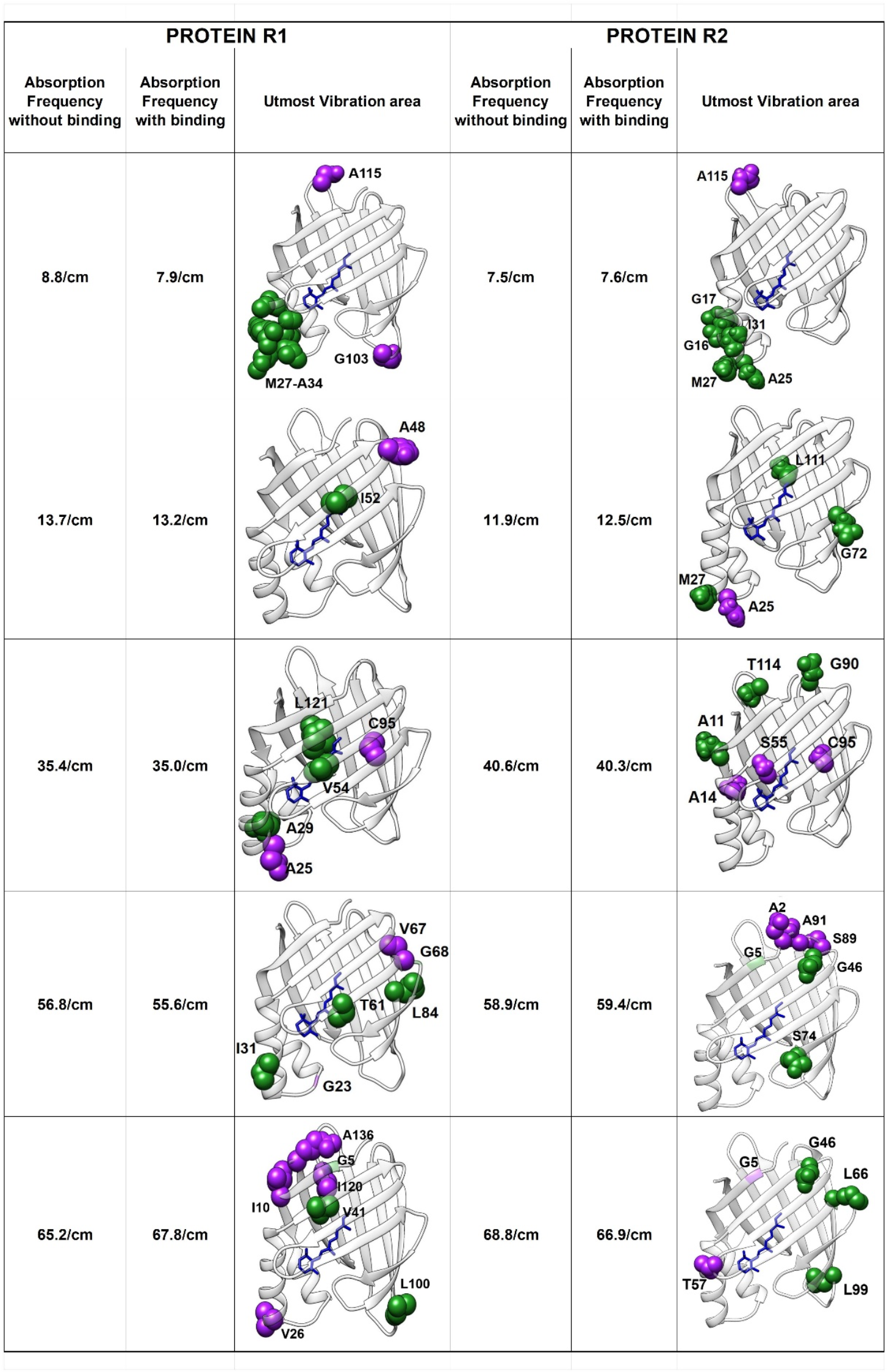
Vibration region analysis with and without retinal binding. The purple spherical area represents the vibration regions without retinal binding, the green globular region represents vibration regions with retinal binding, and the structure in blue stick is retinal.

As can be seen from **Figure 6**, for mutant R1 the vibration regions of the protein without retinal are usually located far away from the cavity where the retinal can be bound. With binding to retinal, the vibration regions are shifted towards the retinal. Taking the lowest vibration frequency as an example, when it’s not bound to retinal, R1 protein has a terahertz absorption frequency of 8.8cm^−1^, and the residues that vibrate strongly are A115 and G103, which are both at the loop regions at the corners of the structure. With binding to retinal, the vibration regions are shifted to the regions from R17 to A34. By measuring the distances between these residues and retinal, we found that they are in a distance less than 5Å from retinal, indicating that interaction between the retinal and protein affects the vibration mode. Accordingly, the absorption frequency of terahertz is shifted to 7.92cm^−1^. Similarly, at the high-frequency domain of protein R1 such as 56.76cm^−1^, the vibration regions are around V67 and G23 in the loop region. With binding to retinal, the vibration regions are shifted to residues I31, T61, and L84, which are in the secondary structures closer to the retinal. For mutant R2, we observed similar behaviors except for the eigenfrequencies around 40cm^−1^ and 68 cm^−1^. The above evidence suggests that binding to retinal introduced interactions between the residues around the cavity and the retinal, which affects the vibration modes.

Clearly, the exceptional behavior at frequencies 40/cm and 68 /cm is due to the effect of the mutation A32W. As we have already seen in Figure 5, the mutation A32W introduces small changes in the vibration regions in the low-frequency domain, but relatively large changes in the middle and high frequency domain. When the alanine 32 is mutated to tryptophan, the π-stacking interaction is introduced between the benzyl-like sidechain of tryptophan and the β-ionone of the retinal, which stabilized the retinal and its neighboring regions^[21]^. As a result, those regions become hard to vibrate, leading to the different behaviors from R1 at frequencies 40/cm and 68 /cm.

On the effects of mutation A32W without retinal binding, we observe very small changes in the vibration regions in the low-frequency domain, but relatively large changes in the middle and high frequency domain, as we have already seen in **Figure 5**. However, we did not observe significant shift of the vibration regions towards the retinal. Hence, we infer that the mutation A32W mainly introduce slight changes in the global vibrations when the mimics are not bound to retinal.

### 3: Comparison of the impact of retinal binding on THz absorption spectra in different mutants

In our study, mutants R1 and R2 differs only in one mutation A32W. Clearly, the single mutation A32W did not change the overall spectra as we see in **Figure 4**. However, when we study how the retinal binding affects THz absorption, such a mutation makes significant difference. We characterized the impact of retinal binding by taking the difference in the terahertz spectra between the same rhodopsin mimic without and with retinal binding. In order to show that the observed phenomenon is robust and general, we calculated the terahertz spectra difference mentioned above for all the samples with different thickness.

**In Figure 7a**, we show the difference of the terahertz absorption spectra between without and with retinal binding for R1 and R2, denoted as ΔR1 and ΔR2, respectively. We can see that ΔR1 and ΔR2 do not show significant divergence until the frequency is larger than 30cm^−1^. When the frequency is larger than 30 cm^−1^, ΔR1 are always positive while ΔR2 are always negative for all three thicknesses. This suggests that binding to retinal weakens the terahertz absorption of mutant R1, but enhances the absorption of R2 protein in the mid-high frequency domain (30∼70 cm^−^ _1_). Considering the sequence difference between R1 and R2, we conclude that the single mutation A32W leads to the different effects of retinal binding on the terahertz absorption in the mid-high frequency domain. In the low frequency region (0∼30 cm^−^ _1_), the mutation A32W does not show any significant effects, which is consistent with our observation in **Figure 5** and **Figure 6**. In **Figure 7b**, we show the difference of the theoretically calculated terahertz absorption spectra between without and with retinal binding, which shows similar properties to **Figure 7a** although there is some detailed mismatch at round frequency 55 cm^−1^.

**Figure 7:**
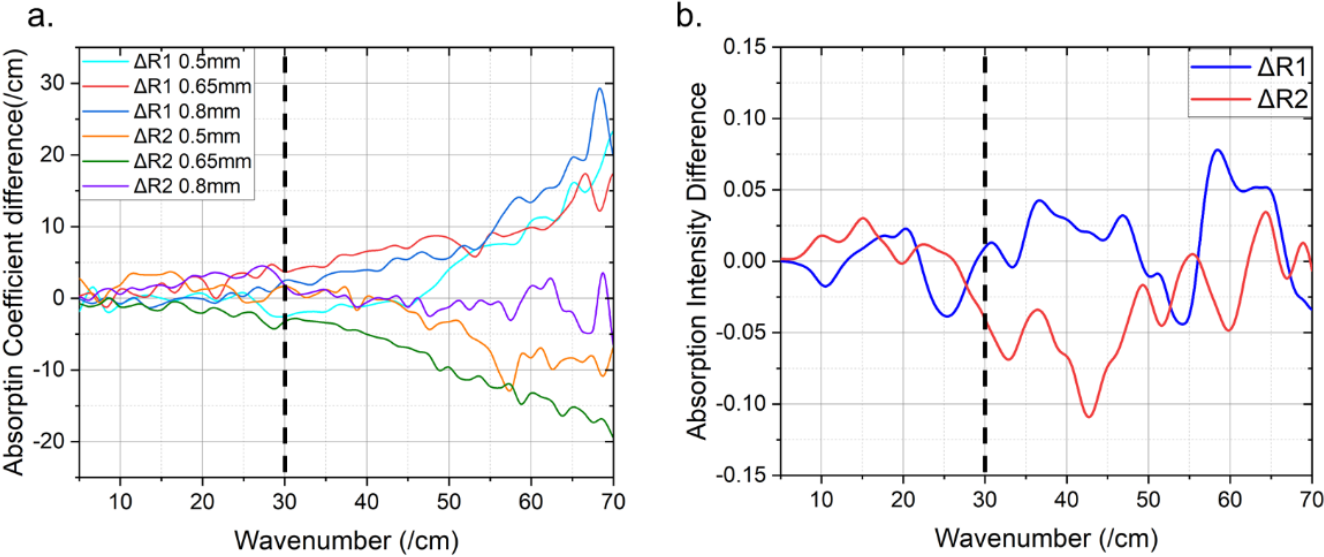
The difference of the Terahertz absorption spectra of the two proteins between without binding and with binding to retinal. a. The experimental result of the difference of the terahertz absorption spectra for R1 and R2 samples with different thicknesses. b. The calculated difference of the terahertz absorption intensity for both proteins using NMA.

We note that the mutation A32W has been proven to have significant impacts on the UV/VS absorption spectra, where the absorption peak of R2 bound with retinal is significantly blue shifted relative to R1^[29]^. In our current study, by comparing the terahertz absorption difference between the cases with and without retinal binding, we show that the mutation A32W also significantly affects the impacts of retinal binding in the absorption in the 30∼70 cm^−1^ terahertz domain. However, the underlying mechanism for this effect still remains to be further investigated.

## Conclusion

In this paper, the effects of retinal binding and the single-point mutation on THZ absorption spectra were investigated by terahertz time-domain spectroscopy and NMA techniques. Our result indicates that the retinal binding exhibited a significant effect on the THz absorption spectra as well as the vibration modes. Specifically, the binding of the retinal causes the shift of the vibration regions towards the retinal. Additionally, by comparing the spectra between two mutants, we found that the single mutation A32W significantly change the effects of the retinal binding on the terahertz absorption. For mutant R1 with residue 32 being alanine, the binding to retinal decreases the terahertz absorption; In contrast, for mutant R2 with residue 32 being tryptophan, the binding of retinal enhances the terahertz absorption. Although the detailed mechanism still remains to be investigated, such an observation indicates A32W has an important role in modulating the vibration modes when the rhodopsin mimic binding to retinal.

By investigating the low-frequency terahertz absorption spectra in rhodopsin mimics as well as their vibration modes, we can gain additional insights into the dynamics of rhodopsin mimics binding chromophores. Such information could be helpful for better understanding the photoreaction process of the rhodopsin mimics and designing new photosensitive proteins.

