## Supplemental Figure for "Influences of single mutation and retinal binding on the THz absorption spectra of CRABP-II based rhodopsin mimics"

**This PDF file includes:**
Supporting text.
Figures S1 to S6;

### Supporting Information Text

#### Refractive index and absorption coefficient calculations

According to the method proposed by Duvillaret et al. in 1996 to obtain the optical parameters of THZ wave of samples, for thick samples, the main peak $E_{sam}(t)$ can be well separated from the first echo peak $E_{echo1}(t)$ in the time-domain spectra, so the required optical parameters can be obtained only with the main peak information.

In the actual experiment, the time-domain spectrum $E_{ref}(t)$ of terahertz wave $E_0$ after passing through the reference nitrogen environment is obtained as the reference signal. Under the same environment and temperature, the time domain spectrum $E_{sam}(t)$ after passing through the sample is measured as the sample signal. They are then Fourier transformed to obtain their frequency spectra $E_{ref}(\omega)$ and $E_{sam}(\omega)$ . And since most substances that absorb terahertz waves meet the extinction coefficient $k_s \ll 1$ and Lambert's Law. The following relation can be obtained by simplification:

$$\frac{E_{sam}(\omega)}{E_{ref}(\omega)} = \frac{4n_s(\omega)}{[1+n_s(\omega)]^2} \cdot exp\left[-i\left(n_s(\omega) - i\frac{\alpha_s(\omega)c}{2\omega} - 1\right)\frac{\omega d}{c}\right]$$

$$= \frac{4n_s(\omega)}{[1+n_s(\omega)]^2} \cdot exp\,[-\frac{\alpha_s(\omega)d}{2}] \cdot exp\left[-i(n_s(\omega) - 1)\frac{\omega d}{c}\right]$$

$$= A(\omega) \cdot \exp\,[i\varphi(\omega)]$$

$A(\omega)$ represents the ratio of the amplitude of the signal in the frequency domain between the protein sample and the nitrogen environment, and $\varphi(\omega)$ represents the phase difference of the signal in the frequency domain between the protein sample and the nitrogen environment.

Therefore, it can be obtained from the above formula:

$$A(\omega) = \frac{4n_s(\omega)}{[1 + n_s(\omega)]^2} \cdot exp\,[-\frac{\alpha_s(\omega)d}{2}]$$

$$\varphi(\omega) = -(n_s(\omega) - 1)\frac{\omega d}{c}$$

Finally, the optical parameters of the protein sample in the terahertz band can be obtained. The refractive index $n_s(\omega)$ and the absorption coefficient $\alpha_s(\omega)$ can be expressed as, respectively:

$$n_s(\omega) = 1 - \frac{c}{\omega d}\varphi(\omega)$$

$$\alpha_s(\omega) = -\frac{2}{d} ln\frac{A(\omega)[1 + n_s(\omega)]^2}{4n_s(\omega)}$$

where $n_s(\omega)$ is the refractive index of the sample. $\alpha_s(\omega)$ is the absorption coefficient of the sample in the terahertz band. φ is the phase difference between the sample signal and the reference signal after the Fourier transform. $c$ is the propagation velocity of the THz wave in a nitrogen environment, ω is the angular frequency, and $d$ is the sample thickness.

#### Figure S1

a.

Protein Sequence

| | 10 | 20 | 30 | 40 | 50 |
| --- | --- | --- | --- | --- | --- |
| R1 | PNFSGNWKII | RSENFEELLK | VLGVNVMLRK | IAVAAASKYA | VEIKQEGDTF |
| R2 | PNFSGNWKII | RSENFEELLK | VLGVNVMLRK | IWVAAASKYA | VEIKQEGDTF |

| | 60 | 70 | 80 | 90 | 100 |
| --- | --- | --- | --- | --- | --- |
| R1 | YIKVSTTVYT | TEINFKVGEE | FEEQTVDGRP | CKSLVKWESE | NKMVCEQKLL |
| R2 | YIKVSTTVYT | TEINFKVGEE | FEEQTVDGRP | CKSLVKWESE | NKMVCEQKLL |

| | 110 | 120 | 130 | 137 |
| --- | --- | --- | --- | --- |
| R1 | KGEGPKTSWT | KELTNDGELI | LTMTADDVVC | TQVFVRE |
| R2 | KGEGPKTSWT | KELTNDGELI | LTMTADDVVC | TQVFVRE |

b.

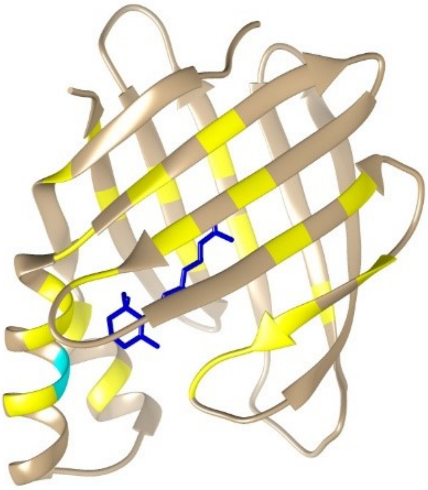

**Figure S1: a. Aligned sequence of R1 and R2.** The mutation A32W are highlight in yellow. The residues that are within 5Å from retinal with binding are colored in red font. **b. The 3D structure of the rhodopsin mimics.** The structure in dark blue is the retinal, the bright yellow part is the residues less than 5Å away from the retinal after binding with the protein, and the mutation A32W is colored in cyan. Notably, the A32W is also within 5Å from the retinal.

**Figure S2**

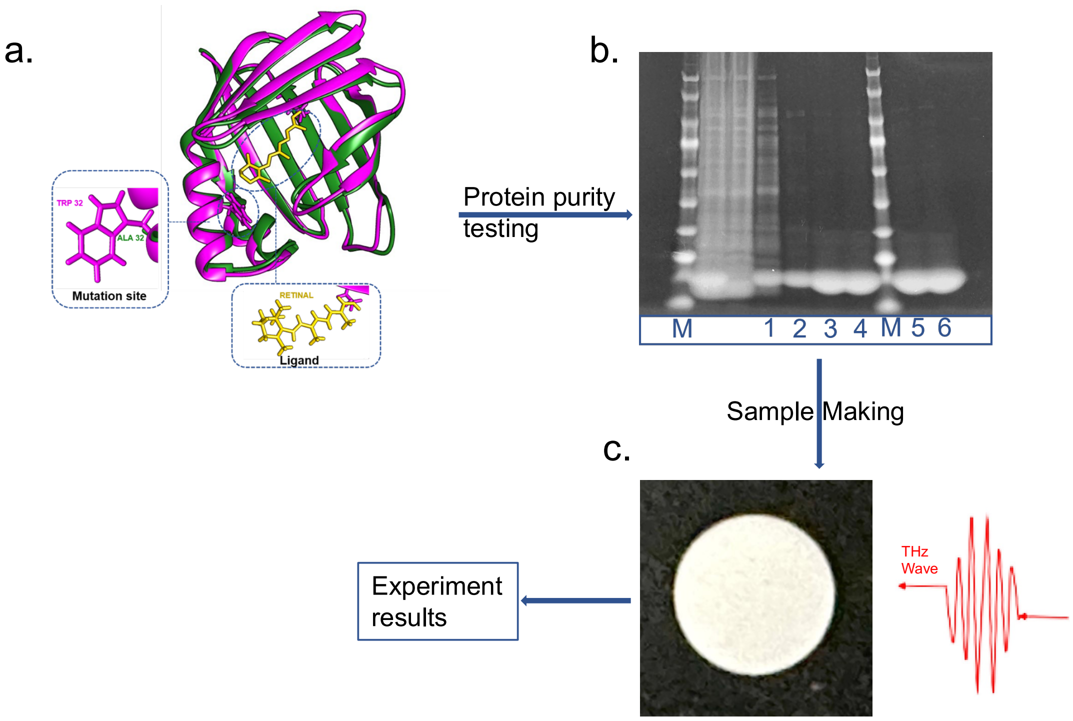


**Figure S2: Schematic diagram of the experiment.** a. Schematic diagram of rhodopsin mimic structure. The green part is the structure of mutant R1, and the purple part is the structure of mutant R2. R1 and R2 mutants differ only in one A32W residue. Among them, the residue includes alanine in R1 and tryptophan in R2. Both mutants have a packet, with the all-trans-retinal chromophore (SI Fig.1 yellow structure) bound to the pocket. The yellow part is the ligand for two proteins: all-trans retinal. b. Schematic diagram of sample purity detected by electrophoresis, where bands 3 and 5 are purified R1 protein bands and 4 and 6 are purified R2 protein bands, both of which meet the experimental requirements. c. Schematic diagram of the terahertz absorption measurement on the solid tablet sample.

### Figure S3

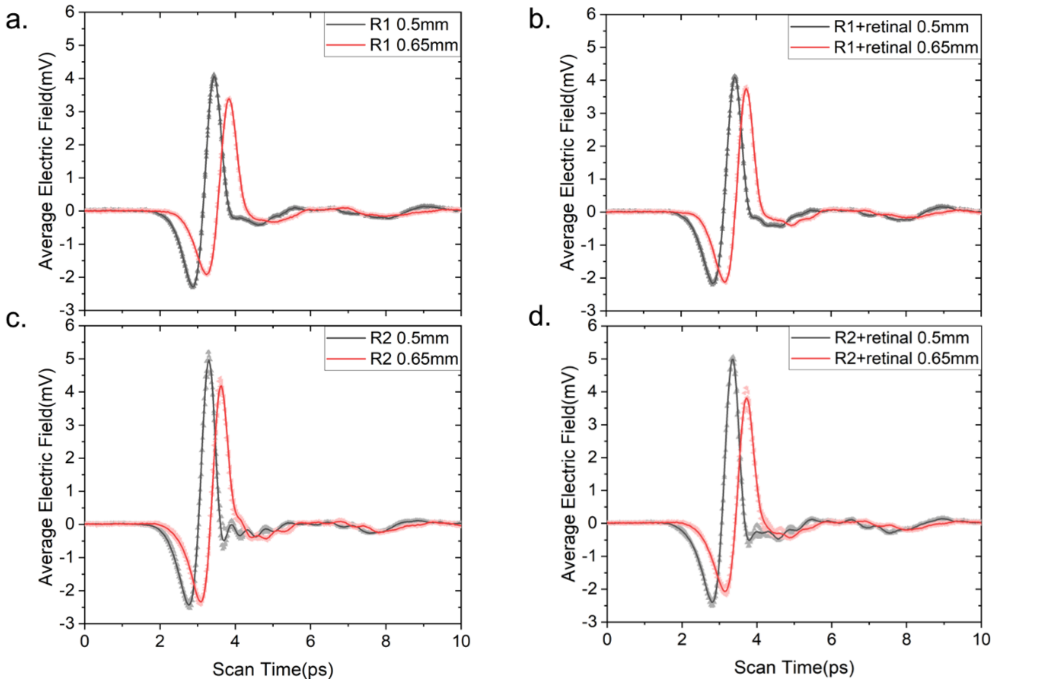


**Figure S3: Terahertz time-domain spectra of rhodopsin mimics with thickness of 0.5mm and 0.65mm.** a. Terahertz time domain spectra of R1. b. Terahertz time-domain spectra of R1 binding with retinal. c. Terahertz time domain spectra of R2. d. Terahertz time domain spectra of R2 binding with retinal.

#### Figure S4

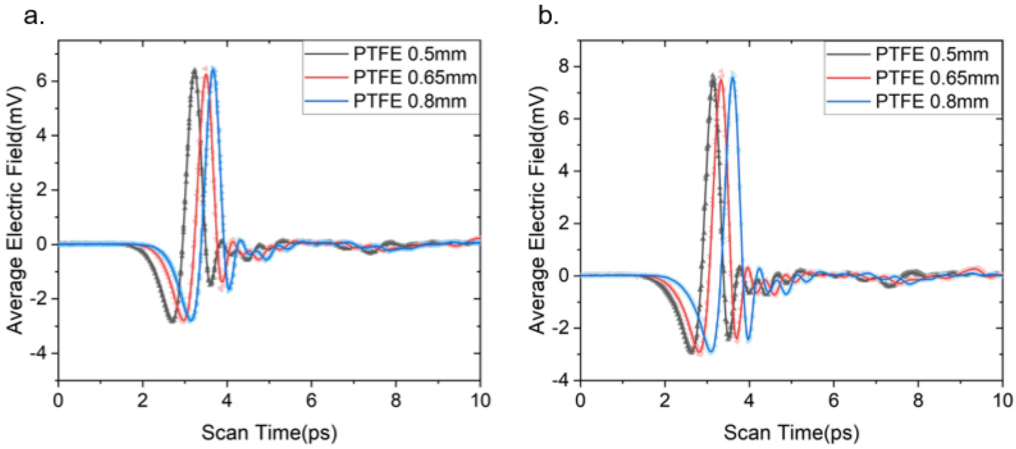


**Figure S4: Terahertz time-domain spectra of the reference PTFE of different thicknesses.** a. terahertz time-domain spectrogram of the reference material PTFE used in the experiment on R1. b. terahertz time-domain spectrogram of the reference material PTFE used in the experiment on R2.

#### Figure S5

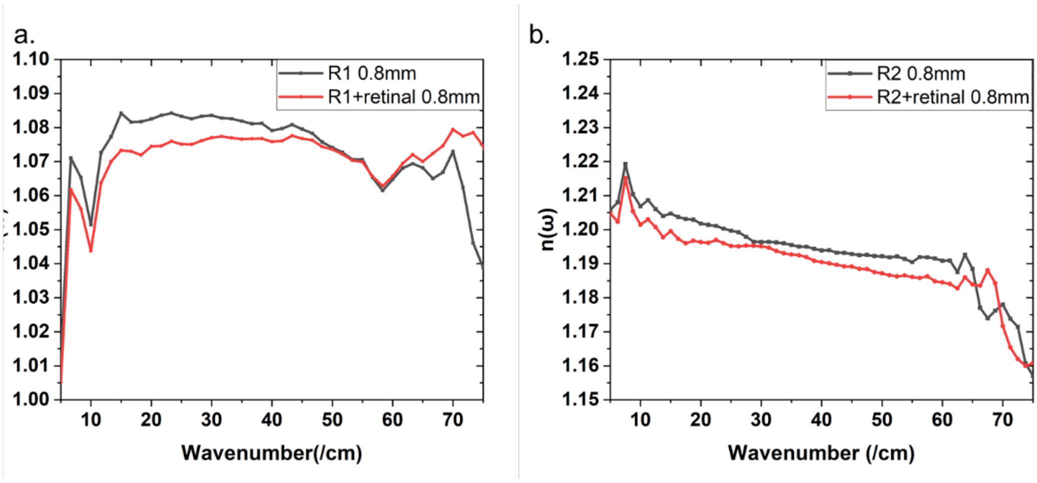


**Figure S5: Refractive index of rhodopsin mimics with sample thickness 0.8 mm in the terahertz band.**

**Figure S6**

**a:**

| PROTEIN R1 | | | |
|---|---|---|---|
| Absorption Frequency without Binding | Utmost Vibration area (Directions included) | Absorption Frequency with Binding | Utmost Vibration area (Directions included) |
| 8.8/cm | 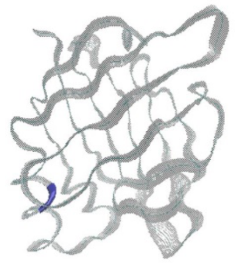 | 7.9/cm | 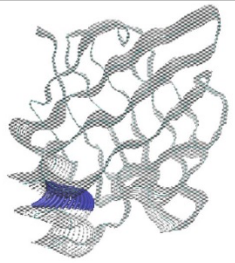 |
| 13.7/cm | 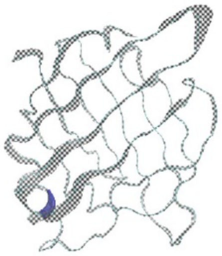 | 13.2/cm | 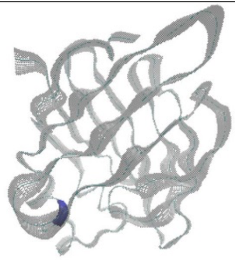 |
| 35.4/cm | 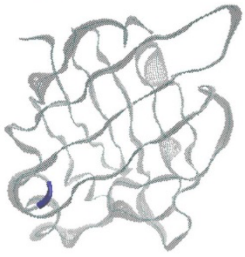 | 35.0/cm | 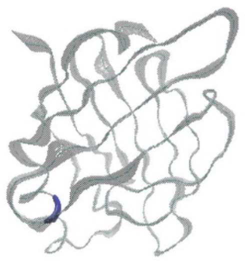 |
| 56.8/cm | 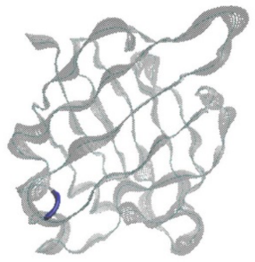 | 55.6/cm | 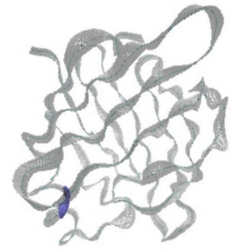 |
| 65.2/cm | 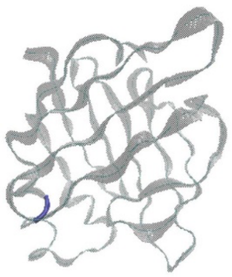 | 67.8/cm | 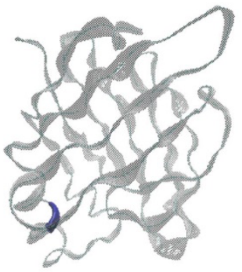 |

**b:**

| PROTEIN R2 | | | |
|---|---|---|---|
| Absorption Frequency without Binding | Utmost Vibration area (Directions included) | Absorption Frequency with Binding | Utmost Vibration area (Directions included) |
| 7.5/cm | 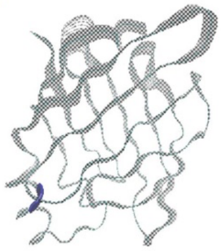 | 7.6/cm | 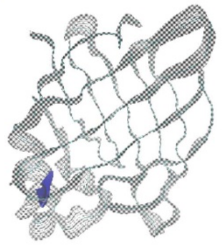 |
| 11.9/cm | 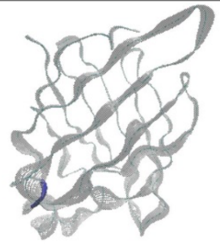 | 12.5/cm | 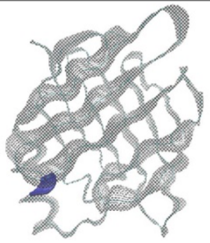 |
| 40.6/cm | 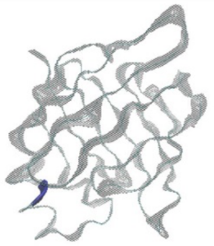 | 40.3/cm | 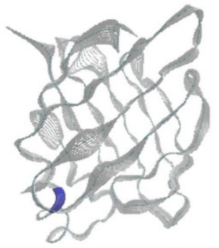 |
| 58.9/cm | 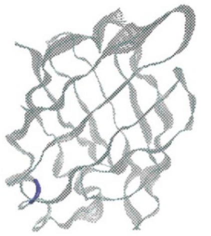 | 59.4/cm | 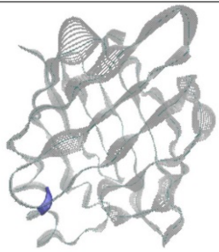 |
| 68.8/cm | 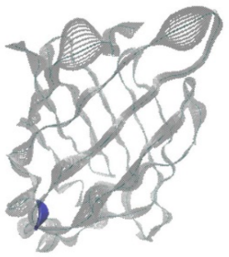 | 66.9/cm | 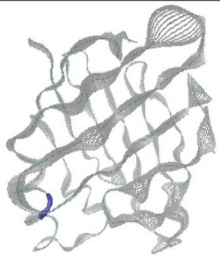 |

**Figure S6: a: Vibration direction of R1 protein at different eigenfrequencies when it binds with retinal and when it is not bound to retinal. b: Vibration direction of R1 protein at different eigenfrequencies when it binds with retinal and when it is not bound to retinal.** (The gray part represents the protein vibration trajectory, while the dark blue part represents the mutation sites of two proteins).
